# Conserved and repetitive motifs in an intrinsically disordered protein drive α-carboxysome assembly

**DOI:** 10.1101/2023.07.08.548221

**Authors:** Julia B. Turnšek, Luke M. Oltrogge, David F. Savage

## Abstract

All photosynthetic bacteria and some chemoautotrophic bacteria fix CO_2_ into sugars in specialized proteinaceous compartments called carboxysomes. Carboxysomes enclose the enzymes Rubisco and carbonic anhydrase inside a layer of shell proteins to increase the CO_2_ concentration for efficient carbon fixation by Rubisco. In the α-carboxysome lineage, a disordered and highly repetitive protein named CsoS2 is essential for carboxysome formation and function. Without it, the bacteria are unable to fix enough carbon to grow in air. How a protein lacking structure serves as the architectural scaffold for such a vital cellular compartment remains unknown. In this study, we identify key residues in CsoS2 that are necessary for building functional α-carboxysomes *in vivo*. These highly conserved and repetitive residues, VTG and Y, contribute to the interaction between CsoS2 and shell proteins. We also demonstrate *in vitro* reconstitution of the α-carboxysome into spherical condensates with CsoS2, Rubisco, and shell proteins, and show the utility of reconstitution as a biochemical tool to study carboxysome biogenesis. The precise self-assembly of thousands of proteins is crucial for carboxysome formation, and understanding this process could enable their use in alternative biological hosts or industrial processes as effective tools to fix carbon.

## Introduction

Carboxysomes are proteinaceous cellular microcompartments that are the metabolic centerpieces of the bacterial CO_2_ concentrating mechanism. Each structure is >100 nm in diameter and encloses the enzymes carbonic anhydrase and Rubisco in an icosahedral-like shell, raising the lumenal CO_2_ concentration and driving Rubisco to operate at its maximum rate and specificity (1–3). There are two carboxysomal lineages that evolved convergently: α-carboxysomes, which emerged in proteobacteria and were horizontally transferred to α-cyanobacteria, and β-carboxysomes, which originated in β-cyanobacteria (4). In this work we focus on the α-carboxysomal lineage, using the proteobacterium *Halothiobacillus neapolitanus* as our model system (5–8).

All carboxysomes require five essential protein components: Rubisco, carbonic anhydrase, hexameric shell proteins, pentameric shell proteins, and a scaffold protein. Much is known about how the enzymatic and shell proteins function in the metabolism and structure of the carboxysome (3, 4, 9, 10), as well as how the β-carboxysome scaffolding protein CcmM drives carboxysome biogenesis within the β-lineage (11–13). Although both carboxysome lineages contain scaffolding proteins, these proteins are related in function alone; they have no sequence or structural similarity. In contrast to the β-lineage, how the α-carboxysome scaffolding protein directs α-carboxysome assembly is far less understood.

The α-carboxysome scaffolding protein, CsoS2, is highly conserved and absolutely required. CsoS2 knockouts cannot produce carboxysomes, rendering the organism incapable of growing at atmospheric CO_2_ levels (0.04% CO_2_) (Fig. S1) (14, 15). CsoS2 is a large ∼90 kDa protein with three distinct domains (Fig. 1A) (16). The N-terminal domain (NTD) contains 4 alpha-helical repeats that bind Rubisco in a low-affinity and multivalent manner (17). The middle region (MR) has 7 distinct repeats, followed by the C-terminal domain (CTD) with 2 repeats and a highly conserved C-terminal peptide (CTP).

**Figure 1.**
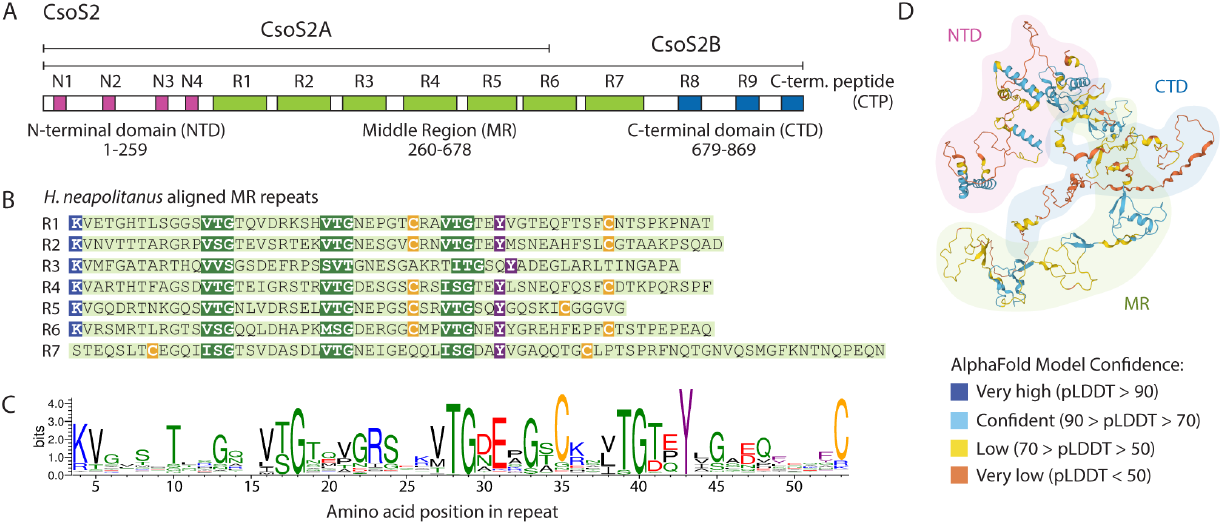
CsoS2 middle region (MR) contains highly conserved and repeated motifs with no known function. (A) Domain architecture of CsoS2. Repeats within domains are indicated by colored blocks. Amino acid numbering is specific to *H. neapolitanus* CsoS2. (B) Alignment of all MR repeats in *H. neapolitanus* CsoS2, with highly conserved motifs highlighted. (C) Sequence logo of the MR repeat generated from an alignment of 1662 MR repeats identified across 272 dereplicated CsoS2 sequences. Blue is basic, red is acidic, green is polar/small, black is hydrophobic, yellow is cysteine, purple is aromatic. (D) AlphaFold model of *H. neapolitanus* CsoS2 (UniProt O85041). pLDDT is AlphaFold’s per-residue confidence score, which scales from 0 to100.

The CTD of CsoS2 binds to shell proteins, and has been successfully used as an encapsulation peptide for heterologous cargo (14, 18, 19). Recently, Ni and Jiang et al. obtained a cryo-EM structure of synthetic mini-carboxysomes with resolved CTD density, showing how it spans across shell-shell interfaces like a staple, reinforcing shell interactions as well as overall mini-carboxysome curvature and T number (20). Interestingly, CsoS2 has a short (CsoS2A) and long (CsoS2B) form produced by a ribosomal frameshifting site in the 6th MR repeat, effectively cutting off the CTD in the short form (21). While the long form is essential for carboxysome formation, the short form is not, further establishing the critical role of the CTD (22). In *H. neapolitanus* both CsoS2A and B are found in equimolar ratios within the carboxysome (14, 23).

Less well studied is the MR domain, which makes up almost 50% of the CsoS2 sequence. The 7 MR repeats have a number of intriguing highly conserved residues and sequence motifs. These were described previously (14, 24) and remain prominent in an up-to-date consensus sequence compiled from 272 de-replicated CsoS2 sequences in which each MR repeat was classified, extracted, and re-aligned against all other individual repeats. Four residues and motifs stand out in particular: (1) (V/I)(T/S)G triplets spaced ∼8-11 amino acids apart (hereafter VTG repeats), (2) cysteine pairs, (3) a highly conserved lysine, and (4) a highly conserved tyrosine (Fig. 1B and C).

In addition to this repeated motif structure, CsoS2 is an intrinsically disordered protein (IDP) as identified by computational disorder predictors and corroborated by circular dichroism spectroscopy (17). CsoS2, like most IDPs, stymies AI structure prediction programs -AlphaFold yields a disordered coil and low confidence scores (Fig. 1D) (25). While it accurately depicted the known NTD alpha helices, it had medium to poor performance predicting the CTD structure, and the MR has never been resolved in either cryoEM or cryo-electron tomography (20, 26, 27).

How an intrinsically disordered protein directs the assembly of the α-carboxysome, and the role of the MR’s highly conserved residues, remains unknown. In this study, we show that some, but not all, of these residues are essential for the growth of *H. neapolitanus* in air. These residues bind to shell proteins in a weak yet highly multivalent fashion and also facilitate the formation of biological condensates when mixed with shell *in vitro*, which may mimic *in vivo* assembly. The *in vitro* condensates show variable liquid properties between shell and CsoS2 under reducing conditions, suggesting a multi-layered assembly strategy optimized for shell localization to the exterior of the carboxysome while minimizing the escape of CsoS2 and Rubisco throughout carboxysome biogenesis.

## Results

### Some, but not all, highly conserved motifs in the CsoS2 Middle Region are essential for carboxysome formation

To probe the function of the highly conserved motifs in the MR repeats, we mutated these residues and assayed the growth of *H. neapolitanus* in air. *H. neapolitanus* needs carboxysomes to grow in air (0.04% CO_2_), but not at higher CO_2_ concentrations, enabling a selection system for deleterious CsoS2 variants. Because the MR has 7 repeats and binding may display complicated behavior, a series of mutants was generated until the entire MR was disrupted (Fig. S2). VTGs were mutated to AAA, Y to A, K to A, and C to S. All strains were generated by knocking out the genomic copy of CsoS2 and re-inserting a complement or mutated copy into a neutral site on the genome (Fig. S1). All strains expressed similar amounts of CsoS2 (Fig. S3), though it should be noted that only CsoS2B was detected; it is likely that expression from the neutral site instead of the native operon reduced ribosomal frameshifting responsible for the production of non-essential CsoS2A.

CsoS2 cysteine and lysine deletion strains showed no loss of growth in air (Fig. 2A, B and E, F), while VTG and tyrosine mutants showed a dramatic loss of growth (Fig. 2C, D and E, F). Growth was dependent on the number of VTG or Y motifs mutated, showing greater attenuation with more mutated repeats. No loss of growth among the cysteine mutants was quite unexpected due to the purported role of redox in carboxysome formation (12, 16) and the seemingly obvious disulfide-bonding function of the conserved cysteine pairs. Despite the growth of CsoS2 cysteine mutant strains of *H. neapolitanus*, CsoS2 cysteine mutant carboxysomes could not be purified from *E. coli* (Fig. S4), suggesting that cysteines play a non-essential structural role that strengthens the overall integrity of the complex, but may not be necessary for its assembly or function.

**Figure 2.**
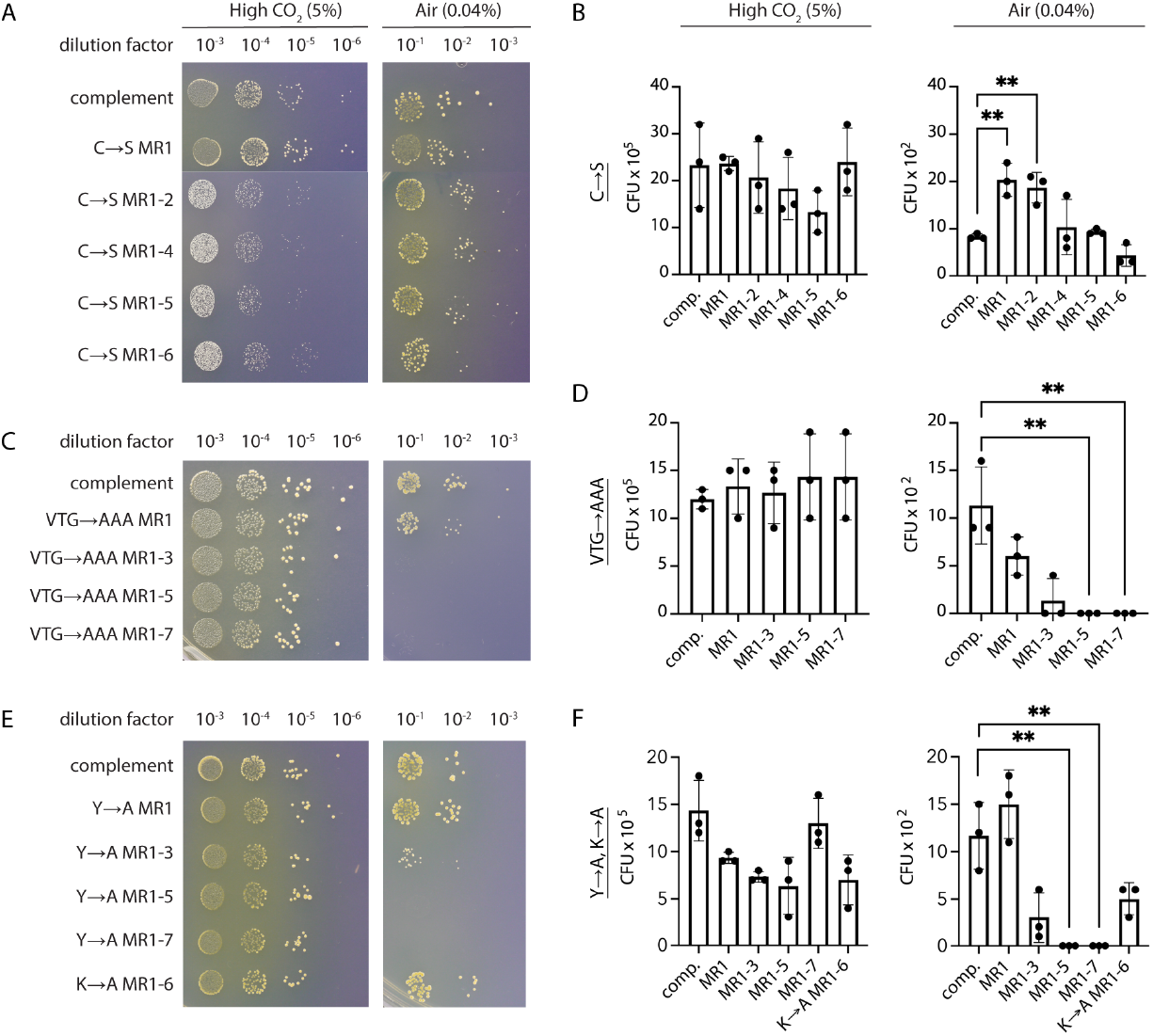
VTG and Y mutations significantly affect growth of *H. neapolitanus* in air. (A, C, E) Representative plates from dilution spotting assays of CsoS2 mutants in *H. neapolitanus*, grown in high CO_2_ (5%) and air (0.04%). (B, D, F) Quantification of spotting assay results. Significance of ** is P ≤ 0.01 in an unpaired t-test. (A, B) C→S mutants, (C, D) VTG→AAA mutants, and (E, F) Y→A and K→A mutants. MR, Middle Region. CFU, colony forming units. Comp, complement.

### The CsoS2 Middle Region binds to shell proteins

Dramatic loss of growth in VTG and Y mutant strains hinted that MR residues form interactions that are essential for carboxysome assembly. Previous studies showed full-length CsoS2 binds to shell protein CsoS1A (14), narrowing down candidate MR interaction partners to either shell proteins and/or CsoS2 itself. To biochemically assess the MR’s binding interactions, we purified full-length CsoS2 and wild-type MR (wtMR) along with VTG and Y mutant variations of wtMR (Fig. 3A). In purifying wtMR, we wanted to identify MR interactions specifically since the NTD was already known to bind Rubisco (17) and CTD to shell (19, 20). Repeat 7 was left out of the wtMR construct because it occurs after the ribosomal slip site in Repeat 6, in an effort to eliminate potential confounding variables between the CsoS2A and CsoS2B isoforms.

**Figure 3.**
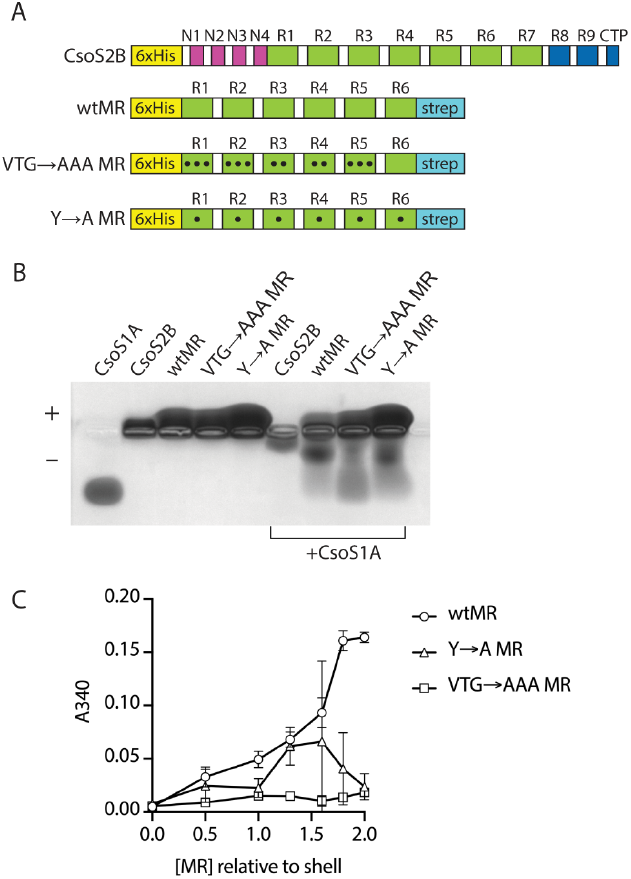
Shell protein CsoS1A binds to the MR of CsoS2, and mutations to VTG and Y residues perturb binding. (A) Purified constructs used in assays; all constructs have the ribosomal slip-site in R6 mutated to only produce the long form. Black circles indicate mutations within a repeat. Cartoons are not to scale with respect to domain sizes. (B) Native agarose protein gel of purified CsoS1A and CsoS2 variants. Quantification of this gel can be found in Fig. S5. (C) Turbidity at 10 minutes of the indicated constructs with defined molar ratios to shell (CsoS1A).

A native agarose electrophoretic mobility shift assay revealed that both CsoS2 and wtMR bind to the hexameric shell protein CsoS1A, and that mutating VTG and Y perturbed this interaction (Fig. 3B and Fig. S5). CsoS1A showed a dramatic shift in mobility when mixed with full-length CsoS2B, and slightly less when mixed with wtMR. Mutating Y led to a subtle but even further decrease in shell mobility, while the mutant VTG construct showed the least binding to shell. We did not evaluate binding to Rubisco or CsoSCA, since it was previously shown that the MR does not interact with either of these proteins (17, 28).

Seeing the binding differences in this qualitative assay, we sought to further investigate the nature of the MR binding interaction. Interactions formed by disordered and/or repetitive proteins can often be monitored by a change in turbidity, which measures the transition from a soluble protein state to phase-separated condensate states. When more wtMR was added to CsoS1A, turbidity increased in a concentration-dependent manner, while no turbidity was observed for each protein alone (Fig. 3C and Fig. S6). The mutant VTG construct had little to no turbidity at any concentration, while the mutant Y construct showed intermediate behavior. Full-length CsoS2 displayed almost 5x more turbidity when mixed with CsoS1A compared to wtMR, reconfirming the robust contribution of the CTD to shell binding (Fig. S6) (18). Taken together with Fig. 3B, these results demonstrate that the VTG and Y residues participate in shell binding to the MR, yet they may contribute to the interaction in distinct ways.

### Formation of phase-separated condensates is dependent on the CsoS2 Middle Region sequence

Following the results of the turbidity assay, we confirmed via fluorescence microscopy that purified CsoS2 and wtMR indeed form phase-separated condensates when mixed with CsoS1A (Fig. 4A). However, when MR with mutated VTG or Y residues was mixed with CsoS1A, no condensates formed. All experiments were performed at 150 mM salt, mimicking typical intracellular conditions, and no condensates appeared when either MR or CsoS1A were observed on their own (Fig. S10). This suggests that these residues may be critical for forming low-affinity, highly multivalent interactions that drive CsoS2 and shell to phase separate during carboxysome biogenesis.

**Figure 4.**
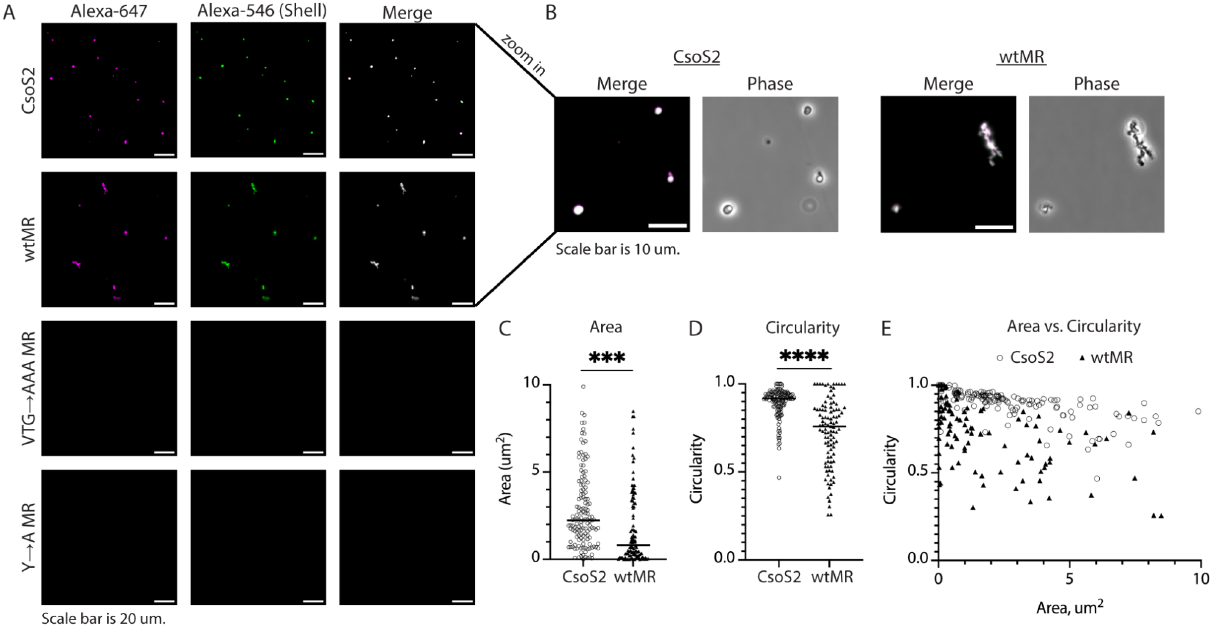
CsoS2 and wtMR form condensates when mixed with CsoS1A, but not when key residues are mutated. (A) Fluorescence microscopy of the indicated CsoS2 / MR protein variants with added CsoS1A, imaged at 30 minutes post mixing. All CsoS2 / MR variants are labeled in pink, CsoS1A is labeled in green, and the merge appears white at equally overlapping intensities. Scale bar is 20 um. (B) Zoom-in of CsoS2 and wtMR droplets shown in (A). Scale bar is 10 um. (C) Comparison of droplet area between CsoS2 and wtMR condensates, measured at 30 minutes. Significance of *** is P ≤ 0.001 in an unpaired t-test. (D) Comparison of droplet circularity between CsoS2 and wtMR condensates, measured at 30 minutes. Circularity is calculated as 4π*area/perimeter^2^, with 1.0 being a perfect circle and lower values indicating increasing shape elongation. Significance of **** is P ≤ 0.0001 in an unpaired t-test. (E) Area vs. Circularity for all measured CsoS2 and wtMR condensates. For (C) and (D), the median is indicated by a black line.

Interestingly, though both CsoS2 and wtMR formed phase separated condensates when mixed with shell, the condensates displayed distinct properties in their size and shape (Fig. 4B). Both condensates showed accumulated growth over 30 minutes (Figs. S7, S8, S9), but tended toward divergent shapes over the same growth period (Fig. S9). CsoS2 condensates were larger and more circular on average, while wtMR condensates were smaller and formed elongated structures (Fig. 4C, D, and E). Condensate shape can be an indicator of a liquid-to-solid phase transition, with liquid droplets often appearing more spherical and solid aggregates appearing more deformed or fibrillar (29, 30). The presence or absence of the CTD in the CsoS2 and wtMR constructs is a proxy for CsoS2A and CsoS2B, suggesting that these two proteins may contribute differently to the physical properties of the nascent carboxysome.

### In vitro carboxysome reconstitution and condensate properties

Since CsoS2 and shell formed phase-separated condensates *in vitro*, we wanted to see if it was possible to fully reconstitute the carboxysome with its three major constituent components: CsoS2, shell, and Rubisco. The NTD of CsoS2 had been previously shown to form phase-separated condensates with Rubisco at low (20 mM) salt (17), but not yet demonstrated with full-length CsoS2 at physiological salt concentrations (150 mM). CsoS2 and Rubisco formed many small condensates at 5 minutes post mixing, but these condensates appeared to dissolve back into the soluble phase over 30 minutes (Fig. S11, S13). In contrast, when CsoS2, Rubisco, and shell (CsoS1A) were mixed, they formed robust condensates that grew significantly in size over 30 minutes (Fig. 5A and Fig. S12, S13).

**Figure 5.**
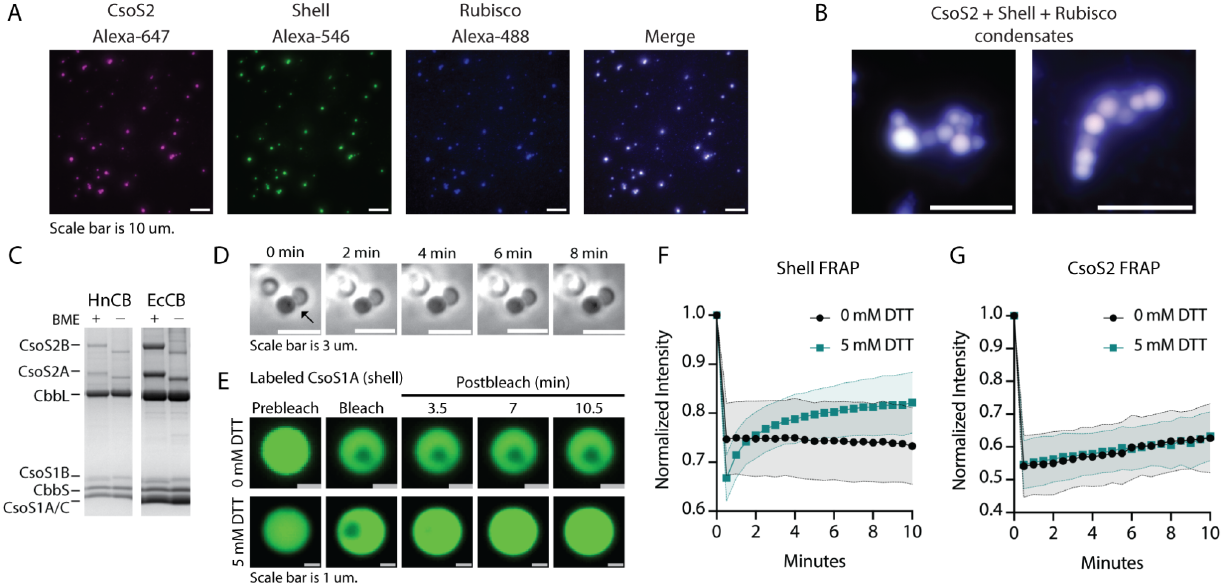
CsoS2, Shell, and Rubisco form condensates with liquid properties that differ in reducing vs. oxidizing conditions. (A) Individual channels of CsoS2 (pink) + shell (green) + Rubisco (blue) condensates, with the merge shown in white. Image was taken 30 minutes post mixing. (B) Zoomed-in examples of merged CsoS2 + shell + Rubisco condensates; scale bar is 10 um. (C) PAGE gel of carboxysomes (CBs) purified from *H. neapolitanus* (HnCB) and *E. coli* (EcCB) with or without β-mercaptoethanol (BME). CsoS2A and B show distinct downshifts under oxidizing conditions. (D) Phase contrast of CsoS2 + shell + Rubisco condensates that do not merge over 8 minutes. (E) Example droplets from FRAP showing labeled CsoS1A with and without 5 mM DTT. (F) CsoS1A FRAP. (G) CsoS2 FRAP. (F, G) bleach occurs at ∼30 seconds; black circles, 0 mM DTT; teal squares 5 mM DTT; normalized intensity, see Experimental Procedures.

To better understand the 3-component condensate formation, CsoS2 and Rubisco were mixed and observed in a gasket on a microscope slide before adding shell at 10 minutes (Fig. S14). Condensates appeared within 30 seconds after the addition of shell. Larger droplets settled onto the focused plane of the slide over time, suggesting that droplet growth may occur via accretion of individual soluble components. There was also no significant difference in the average number of CsoS2-Rubisco-shell condensates per micrograph between 5 minutes and 30 minutes, further supporting a growth mechanism triggered and driven by the presence of shell (Fig. S13).

Interestingly, condensates were often observed to adhere next to one another without merging over time (Fig. 5B and D), a behavior that implied a more gel-like than liquid-like state (29). Protein liquidity in condensates can be sensitive to the solvent chemical environment, including to reducing / oxidizing (redox) conditions (12, 31, 32), which can alter local structure and dynamics via changes to reactive moieties like cysteine side chains.

We found that redox-dependent behavior was present in both intact carboxysomes and in reconstitutions. When carboxysomes, purified from either native or heterologous sources, were analyzed by SDS-PAGE under both reducing and oxidizing conditions, CsoS2A and CsoS2B – and no other constituents – display a marked size shift, running as smaller under oxidizing conditions (Fig. 5C). Notably, this was a change in size from one homogenous species to another, suggesting that CsoS2 may undergo a specific redox-dependent structural change which could affect interactions between CsoS2 and its binding partners. To test this hypothesis, the liquidity of individual components was assessed using fluorescence recovery after photobleaching (FRAP). FRAP of CsoS2-Rubisco-shell condensates revealed that the shell experiences a dramatic difference in mobility depending on the redox conditions; it had no mobility under oxidizing conditions and recovered under reducing conditions (Fig. 5E and F). In contrast, CsoS2 showed nearly identical slow recovery under both oxidizing and reducing conditions (Fig. 5G). These results emphasize that the redox environment can independently modulate the mobility of proteins in carboxysome condensates, selectively tuning condensate properties and providing a window into how *in vivo* carboxysome assembly functions.

## Discussion

Carboxysome assembly spans across length scales, from single amino acid interactions to thousands of proteins organizing themselves into a >300 megadalton compartment. In this work, we dissect each length scale to form a new model of how CsoS2 coordinates assembly of α-carboxysomes. With *in vivo* studies in the native host organism, we identified VTG and Y sequence motifs in the MR as essential for carboxysome assembly. These motifs appear to interact with the shell, an effect which is amplified by high valency across 7 MR repeats. Mutation of key resides *in vivo* and *in vitro* weakened this interaction. We further demonstrate that CsoS2, Rubisco, and shell can be reconstituted *in vitro* into spherical condensates, and that the liquidity of the shell can be tuned by the redox environment.

Overall, VTG and Y motifs contribute to many weak, transient interactions that en masse increase the binding affinity to shell proteins (Fig. 6A). It’s not obvious at first glance how (V/I)(T/S)G motifs facilitate binding to the shell. A small step away to alanine abolished binding, suggesting important contributions from additional alkyl groups as well as the hydrogen bonding interactions from the hydroxyl group. In contrast, there is precedence for the importance of tyrosine residues in intrinsically disordered proteins (IDPs). Tyrosines often participate in pi-pi or cation-pi interactions with other aromatic or charged residues. IDPs with repetitive Y residues such as Fused in Sarcoma (FUS) showed reduced phase separation when greater numbers of Ys were mutated (33–35), similar to the MR.

**Figure 6.**
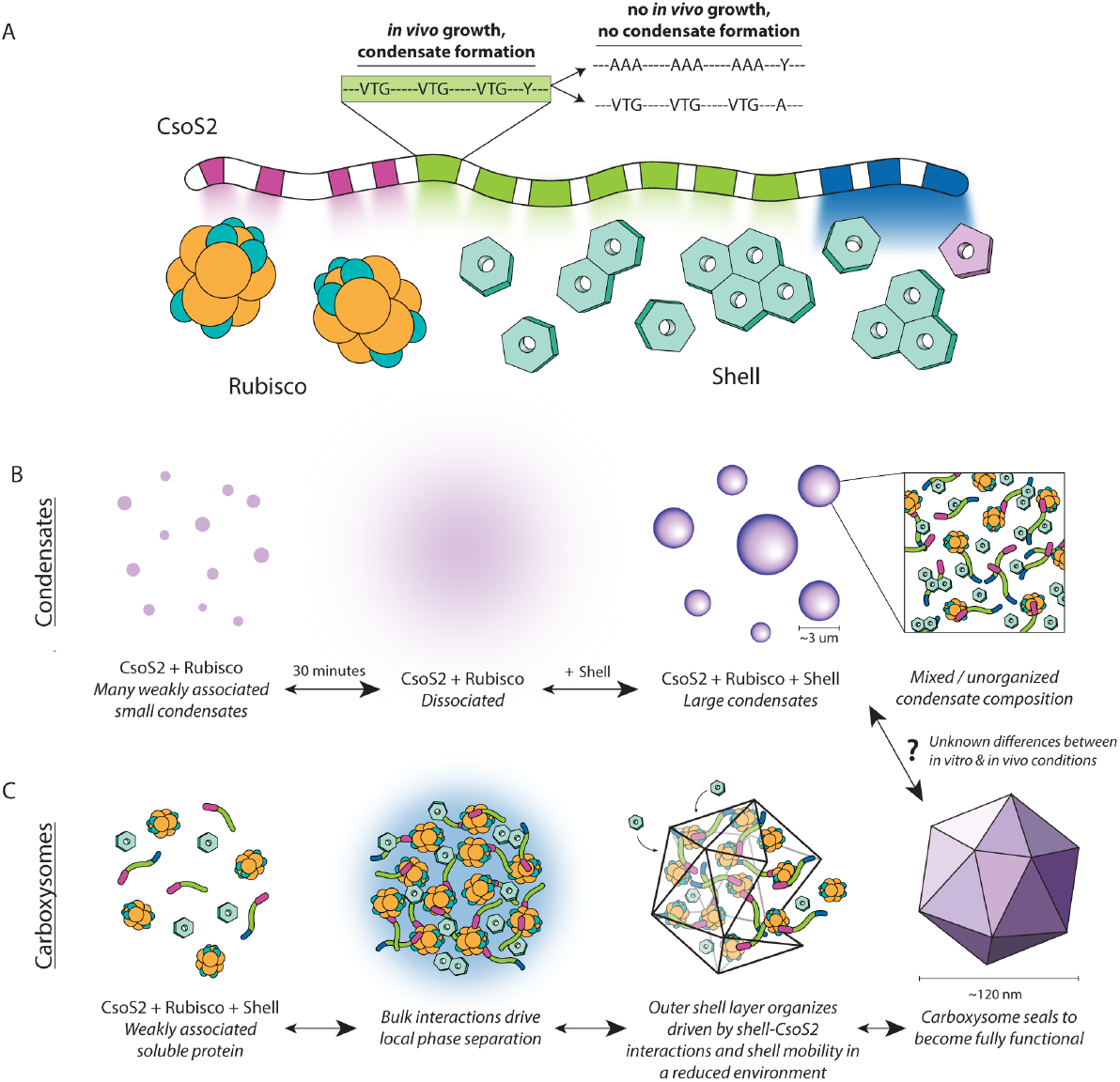
Model of CsoS2 interactions driving condensate and carboxysome assembly. (A) Cartoon model of known interactions between CsoS2 motifs and binding partners, with colored spotlights highlighting the relative strength of each interaction. Pink blocks are NTD repeats, green blocks are MR repeats, and blue blocks are CTD repeats and the CTP. The pullout box shows the sequence residues that are important for *in vivo* growth and condensate formation, and the mutated variants that do not grow or form condensates. (B) Model of condensate formation. Weak associations between CsoS2 and Rubisco tend towards dissolution at equilibrium but addition of shell precipitates large condensates. Condensates are assumed to be a mixture of all three proteins with no clear shell layer. (C) Model of *in vivo* carboxysome assembly, informed by condensate biochemistry. All carboxysome components phase separate locally. Shell-CsoS2 interactions drive assembly and organization, leading to the formation of sealed and functional compartments.

The sticker and spacer model has emerged as a useful framework for understanding protein phase separation. In the model proteins are divided into “sticker” regions responsible for intermolecular interactions and intervening disordered “spacers” (36). The “stickers” may range from single amino acid sidechains (such as the Ys in FUS) all the way to well-defined folded domains such as SSULs in the β-carboxysome scaffold protein CcmM (12). It remains to be seen where the MR lies along this spectrum, that is, whether an MR repeat acts as single binding unit or a collection of short motifs, e.g. VTGs. A recently solved structure of the CTD bound to shell shows a well-conserved yet extended conformation (20). AlphaFold structures of the MR repeats suggest a pseudo-threefold arrangement of the three VTG motifs which occur with highly regular spacing. This structure bears striking shape complementarity to the threeway hexamer junctions and the VTGs could plausibly engage with CsoS1A-His79 as observed for the CTD repeat VTG motifs in the structure of Ni et al. (37). It is possible that the highly conserved tyrosines of the MR repeats act as a stabilizing core for the VTG triads and make the MR repeats into a unitary interactor.

If MR binds to shell proteins, what is its role in the context of CsoS2B, which includes the additional shell binding CTD domain? Valency, sequence, and charge are three key differences. In *H. neapolitanus*, the MR has a valency of 7 repeats, while the CTD only has 2. Although these two repeats possess VTGs, they lack the conserved tyrosines, cysteines, and lysines (Fig. S15), and are also followed by the C-terminal peptide. The CTD has a pI of 9.4, making it positively charged at pH values close to 7 and promoting interaction with the negative shell lumenal interface (18, 20). In contrast, the MR has a pI of 6.2. The roughly 2:1 ratio of MR to CTD (based on the 1:1 ratio of CsoS2A to CsoS2B) additionally amplifies the MR-shell interaction. However, too high of an A:B ratio may be detrimental to carboxysome formation -*in vitro*, MR and shell formed elongated condensates while CsoS2B and shell formed spherical condensates, and *in vivo*, CsoS2A alone is not sufficient to form carboxysomes (21). These differences likely act in concert to give the MR a mode of binding to the shell that is distinct from the CTD. Recent work from Oltrogge et al. proposes that the MR repeats bind areas of less shell curvature, while the CTD favors higher curvature, thus serving to template carboxysome size (37).

The ability to study carboxysome assembly both *in vivo* and in *vitro* has many benefits, though it must be acknowledged that carboxysome condensates are not true carboxysomes. They are thousands of times larger in volume, do not contain all carboxysome components at exact ratios found *in vivo*, and lack the architectural organization of an outer shell layer and icosahedral shape. However, they are extremely useful as a proxy tool to biochemically interrogate protein interactions that are challenging to study *in vivo* (Fig. 6B). From condensates, we learn that CsoS2 and Rubisco are at the edge of solubility and weakly interact at physiological salt concentrations. Addition of shell leads to robust condensate formation, identifying the shell-CsoS2 interaction as the main driver of local phase separation. Notably, there is no evidence of organization in these condensate assays; we assume CsoS2, Rubisco, and shell are homogeneously mixed.

Extrapolating what this tells us about *in vivo* α-carboxysome formation (Fig. 6C), it is known that CsoS2, Rubisco, and shell are transcribed and translated from the same operon in distinct ratios (38, 39). It is thus likely that carboxysome proteins interact immediately during coincident expression and that these initial interactions drive local phase separation. In the reduced cytosol, the shell is more mobile and forms interactions with both CsoS2 and itself both from within the condensate and via outside accretion to organize an outer layer. At a certain volume it becomes thermodynamically favorable for shell proteins to fully encapsulate the carboxysome condensate, blocking additional growth and sealing off a functional carboxysome (40, 41). To kinetically trap carboxysome growth or dissolution at a precise size is perhaps even a role of the shell *in vivo*, in addition to concentrating CO_2_. This is the key step where condensates and carboxysomes differ -it is still unknown what branches the completion of a 150 nm compartment from continued growth to a micron-sized particle. It might simply require tweaking of *in vitro* reaction conditions -salt concentration, protein concentration, molecular crowding, etc. -to tilt the preference towards smaller compartments. Future work will aim to not only establish the precise conditions to form nm-sized carboxysomes *in vitro*, but also to confirm that they can carboxylate CO_2_.

Carboxysomes are a fascinating model system to understand how the coordinated actions of thousands of proteins build an essential cellular structure. Remarkably, the instructions for compartment assembly are encoded solely in the sequences of its constituent proteins. Here we establish that there is a molecular grammar to the CsoS2 MR sequence and how disruption of even a small number of residues diminishes binding to shell and prevents carboxysome formation. This work contributes new motifs to the growing dictionary of known sequence determinants of phase separation and microcompartment formation. Predicting whether a protein will phase separate and form a compartment, along with the conditions that affect this interaction, will continue to be important for informing broader efforts to engineer carboxysomes and other diverse microcompartments in biological systems.

## Acknowledgements

We thank Noam Prywes for thoughtful comments on the manuscript draft. This work was supported by the U. S. Department of Energy, Office of Science, Office of Basic Energy Sciences, Chemical Sciences, Geosciences, & Biosciences Division, Physical Biosciences Program, under Award Number DE-SC0016240 to D.F.S. and D.F.S. is an Investigator of the Howard Hughes Medical Institute.

## Experimental procedures

### CsoS2 MR and CTD consensus sequences

All sequences in IMG matching the CsoS2 pfam (PF12288) were downloaded in May 2020 for a total of 770 sequences. Partial sequences and those with ambiguous residue assignments were discarded, and the set was dereplicated to 95% protein sequence identity using usearch (42). These 272 remaining sequences were analyzed for peptide motifs using the MEME suite (43). The MR and CTD repeat motif positions were identified using MAST for a total of 2190. These repeat sequences were extracted with 15aa of buffer on either side and then all aligned against each other, including both MR and CTD types, using mafft (44). FastTree was used to build a phylogenetic tree of all the repeats which clearly separated into two major clades: one with MR repeats and one with CTD repeats (45). A number of repeats had been misidentified by MAST as evidenced by their membership in the opposing clade. Notable among these is R7 from *H. neapolitanus* which the phylogeny strongly suggests is actually an MR repeat. The sequences belonging to the MR repeat clade (1662) and CTD repeat clade (528) were aligned again with mafft but this time only against members of their respective clades. Weblogo3 was used to create sequence logos for the two repeat classes from these alignments (46).

### *H. neapolitanus* strain generation

Wild type (WT) *Halothiobacillus neapolitanus* is strain c2, ATCC 23641. The *ΔcsoS2* strain was made by homologous recombination of a spectinomycin resistance cassette into the native CsoS2 locus. Complement and mutant strains were generated by homologous recombination of the new CsoS2 sequence into a neutral site on the genome in the *ΔcsoS2* background strain. The insertion region corresponds to bases 2428660 -2429201 on the genome. Plasmids were made by Golden Gate cloning into a neutral site destination vector. The neutral site vector contained the following features from 5’-3’: *H. neapolitanus* upstream homology arm (bases 2428121 -2428660), KanR, LacIQ, pTRC promoter, gene of interest (CsoS2), TrrnB terminator, *H. neapolitanus* downstream homology arm (bases 2429201 -2429703). All sequences contained an intact frame shifting site in CsoS2.

To transform *H. neapolitanus*, 10 ml of DSMZ-68 medium was inoculated per transformation and grown at 30°C and 5% CO_2_. Cells were collected when the pH indicator had turned gray or light yellow (1-2 days of growth). Cells were pelleted at 4000 xG for 10 minutes at 4°C and washed with cold milliQ water twice. After the third spin, cells were resuspended in 50-100 μl cold milliQ water. Cells were mixed with 500 ng of linearized plasmid and placed in cold electrocuvettes, then electroporated at 19 kV/cm, 200 mA, and 25 uF before immediate resuspension in 1 ml cold DSMZ-68 without antibiotic. Recovery occurred during overnight incubation at 30°C and 5% CO_2_ before plating on DSMZ-68 agar plates containing the selection antibiotic. Colonies usually appeared after 3-4 days of growth at 30°C. The genotype was confirmed via colony PCR and sequencing.

### *H. neapolitanus* selection assays

*H. neapolitanus* was inoculated from a colony on a plate into DSMZ-68 medium with the appropriate antibiotic (none for WT, 10 μg/ml spectinomycin for *ΔcsoS2*, and 10 μg/ml spectinomycin + 2 μg/ml kanamycin for complement / mutant strains) and 100 μM IPTG. Colonies were grown 1-2 days in 5% CO_2_ until the medium had turned gray, which corresponded to OD600 ∼0.1-3. Cells were washed with DSMZ-68 without pH indicator added to collect a more accurate OD600. All strains were normalized to OD600 of 0.1 and a 10x dilution series was generated. Strains were plated onto dry DSMZ-68 plates with the appropriate antibiotic and 100 μM IPTG and grown in either 5% CO_2_ or air at 30°C.

### Protein expression and purification

All proteins (His-CsoS2B, 6xHis-wtMR-strep, 6xHis-(VTG→AAA MR)-strep, 6xHis-(Y→A MR)-strep, 6xHis-Rubisco, and strep-CsoS1A) were individually cloned into pET-14-based destination vectors with ColE1 origin, T7 promoter, and carbenicillin resistance. For the VTG→AAA MR construct, all VTG and VSG sites were mutated to AAA in repeats 1-5. For the Y→A MR construct, all Y sites were mutated to A in repeats 1-6. Plasmids were transformed into *E. coli* BL21-AI cells. Cells were grown in LB medium at 37°C with appropriate antibiotic until mid-log phase (OD_600_ of 0.3-0.5), at which point 0.2% L-arabinose was added to induce protein expression and the temperature lowered to 18°C. Cells were grown overnight before pelleting at 5000*g* the next day and freezing at -20°C.

Frozen cell pellets were thawed and resuspended in lysis buffer (50 mM Tris, 300 mM NaCl, 20 mM imidazole, pH 7.5) with the addition of 1 mM phenylmethanesulfonyl fluoride (PMSF), 0.1 μl/ml benzonase, and 0.1 mg/ml lysozyme. Cells were lysed on an Avestin EmulsiFlex-C3 homogenizer and clarified at 27*g* for 45-60 minutes. All subsequent purification steps were performed at room temperature. Supernatant was added to a Ni-Sepharose resin in a gravity column, washed with wash buffer (50 mM HEPES, 300 mM NaCl, 60 mM imidazole, pH 7.5) and eluted with elution buffer (50 mM HEPES, 300 mM NaCl, 300 mM imidazole, pH 7.5). Proteins with a strep tag (His-wtMR-strep, His-(VTG→AAA MR)-strep, His-(Y→A MR)-strep) were further cleaned up on a Strep-Tactin resin on a gravity column. The entire elution was loaded onto the column, washed with wash buffer (50 mM HEPES, 300 mM NaCl) and eluted with elution buffer (50 mM HEPES, 300 mM NaCl, 2.5 mM d-desthiobiotin) before adding 10% glycerol, flash freezing in liquid N_2_, and storing at -80°C.

CsoS2B was purified the same way through the His elution step, then further cleaned up using size exclusion chromatography. Eluted protein was loaded onto a HiPrep 16/60 Sephacryl S-200 HR column equilibrated in 50 mM HEPES, 300 mM NaCl buffer on an Akta Pure chromatography system. Fractions with full-length protein were concentrated on Amicon Ultra 15 Ultracel 30K filters before adding 10% glycerol, flash freezing in liquid N_2_, and storing at -80°C.

Strep-CsoS1A was lysed and clarified the same way as above in 50 mM HEPES, 150 mM NaCl lysis buffer, then purified on a Strep-Tactin resin on a gravity column. Clarified lysate was loaded onto the column and allowed to flow through, followed by a wash step and elution with elution buffer (50 mM HEPES, 150 mM NaCl, 2.5 mM D-Desthiobiotin) before adding 10% glycerol, flash freezing in liquid N_2_, and storing at -80°C.

His-Rubisco was lysed and clarified the same way as above yet with a different lysis buffer (50 mM Tris, 150 mM NaCl, 20 mM imidazole, pH 7.5) and purified on a Ni-Sepharose resin on a gravity column the same way as above. Wash buffer was 50 mM Tris, 150 mM NaCl, 60 mM imidazole, pH 7.5. Elution buffer was 50 mM Tris, 150 mM NaCl, 300 mM imidazole, pH 7.5. Eluted protein was buffer exchanged on a 2 mL Zeba desalting column before adding 10% glycerol, flash freezing in liquid N_2_, and storing at -80°C.

### Carboxysome expression and purification

Carboxysomes were expressed in *E. coli* BW25113 off of the pHnCB10 plasmid (as described in Bonacci et al.(8)) with 500 μM IPTG induction at mid-log phase. Cells were grown overnight at 18°C and pelleted the next day. Carboxysomes were purified as described previously (17). Briefly, cell pellets were lysed using B-PER reagent with the addition of 1 mM PMSF, 0.1 μl/ml benzonase, and 0.1 mg/ml lysozyme. Lysis took place for 45 minutes at room temperature while shaking. Lysate was spun for 20 minutes at 12,000*g*, the supernatant collected, and then spun again for 30 minutes at 40,000*g* and the supernatant discarded. The pellet was resuspended in 200 μl TEMB (10 mM Tris, 10 mM MgCl2, 1 mM EDTA, pH 8.0) on ice with gentle rocking for 1 hour to overnight, with additional resuspension via pipette if needed. The resuspended pellet was clarified for 3 minutes at 1000*g* before loading onto a 5-step sucrose gradient (10, 20, 30, 40, and 50% w/v sucrose in TEMB). Gradients were spun for 15 minutes at 105,000*g* or longer depending on the size of the prep. The gradient was fractionated and analyzed with SDS-PAGE. Fractions containing carboxysomes were pooled and centrifuged for 30-90 minutes at 105,000*g*, resuspended in TEMB, and stored at 4°C. The SDS-PAGE gel in Figure 5C was run with 1% (v/v) β-mercaptoethanol (BME).

### Native agarose protein gel

The gel was made from Tris-acetate-EDTA (TAE) buffer with 1% agarose. Samples were mixed (buffer: 50 mM HEPES, 150 mM NaCl) and cooled to room temperature over 45 minutes before adding native loading dye. Samples were not boiled. 5 μg of protein was loaded into each well for the controls; for mixed samples 5 μg of CsoS1A was added in addition. The gel was run for 60 minutes at 60 volts in native buffer (25 mM Tris, 200 mM Glycine). The gel was stained for 1 hour with Gel Code Blue, then destained with water until most of the stain had dissipated from the background. Gel quantification was done in FIJI.

### Turbidity assays

Protein was thawed to room temperature before mixing. All samples were prepared to a final buffer composition of 50 mM HEPES, 150 mM NaCl. All samples contained 9 μM CsoS1A (except for the 0 μM shell sample). Concentrations of MR variants were: 0, 4.5, 9, 12, 14, 16, and 18 μM. The 0 μM shell control had 18 μM of MR. For CsoS2, concentrations tested were 4.5 and 14 μM. The 0 μM shell control had 14 μM of CsoS2. 40 μl were pipetted into a Nunc 384 well transparent plate and data collected on a Tecan Spark plate reader.

### Condensate microscopy and quantification

Strep-CsoS1A (shell) was labeled with Alexa546 NHS Ester, at a ratio of 2x dye to hexamer. All CsoS2 and MR variants were labeled with Alexa647 NHS Ester, at a ratio of ⅙ dye to monomer. Rubisco was labeled with Alexa488 TFP Ester at a ratio of ⅓ dye to L8S8 hexadecamer. Prior to dyeing, Rubisco was buffer exchanged into 50 mM HEPES, 150 mM NaCl buffer on a Spin-X UF Corning 100K 0.5 ml filter tube. Labeling occurred for 1 hour in the dark at room temperature. Thermo Fluorescent Dye Removal Columns (#22858) were used to wash away the unconjugated dye, using an equal amount of resin to the volume of the sample. Proteins were thawed to room temperature before mixing in a PCR tube in a final buffer concentration of 50 mM HEPES, 150 mM NaCl. All proteins were at a concentration of 10 μM, except for the Rubisco+CsoS2+shell sample, which had 7.9 μM Rubisco, 6.1 μM CsoS2, and 17.5 μM shell. For the gasket experiment (SI Figure 14), 2 μl of shell at 105 μM was added at the 10 minute mark to 10 μM Rubisco and 10 μM CsoS2.

At 5 minutes and 30 minutes, a 1 μl sample was taken from the tube and pipetted onto a microscope slide (VWR micro cover glass 24x60mm No.1) and a coverslip added (VWR micro cover glass 24x30mm No.1). Samples were imaged on a Zeiss Axio Observer Z1 inverted fluorescence microscope at 100x magnification with an oil immersion objective. The gasket in SI Figure 14 is a Coverwell Perfusion Chamber 8x9mm diameter by 0.9mm depth (#622105). The Alexa546 channel appears as green, the Alexa647 channel appears as magenta, and the Alexa488 channel appears as blue.

All images were analyzed in FIJI. The intermodes thresholding algorithm was used to define droplets and make a mask before taking measurements. Condensates under 0.002 um^2^ and over 10 um^2^ were discarded due to false positives of misclassified droplets during the thresholding process.

### FRAP measurements

FRAP experiments were done on a Leica DM6 CFS microscope with a white light laser. Each image was taken as a z-stack. A pre-bleach image was taken, and then droplets were bleached at 499, 557, and 653 nm at 40% laser intensity. A post-bleach timelapse took an image every 30 seconds for 10 minutes. For image analysis, images were first converted into average projections using the LASX microscope software, then further analyzed on FIJI. Drift correction was applied to each channel using StackReg (translation), then the background subtracted using a 50 pixel radius. FRAP measurements were analyzed using the method described in Guillén-Boixet et al. (47).

### Western blots

5 ml of each strain were grown at 30°C and 5% CO_2_ (except for WT which was grown in air) in DSMZ-68 medium with appropriate antibiotic and with or without 100 μM IPTG. At early log phase (indicated by grey or light yellow pH indicator in the medium), cells were pelleted at 4000*g* for 10 minutes, the supernatant discarded, and frozen at -20°C for later analysis. For analysis, cells were thawed with 200 ml B-PER reagent, plus 1 mM PMSF, 0.1 μl/ml benzonase, and 0.1 mg/ml lysozyme (final concentrations). Lysis occurred over 45 minutes at room temperature while shaking. Samples were mixed with loading dye containing BME and boiled for 6 minutes. For the blot in (a), ∼25 μg of protein was loaded per well (+/-3 μg), for (b) 25 μg, and for (c) 50 μg (12.5 μg for WT). Samples were run on a Biorad TGX 4-20% gel for 40 minutes at 180 V. The gel was rinsed in water before transferring onto a PVDF membrane using a Biorad TransBlot Turbo for 10 minutes at 2.5A and 25 V. The membrane was blocked in TBST (50 mM Tris pH 7.5, 150 mM NaCl, 0.1% Tween-20, 5% rehydrated milk) overnight at 4°C while shaking. The next morning the buffer was replaced with 10 ml new TBST (2.5% milk) and primary antibody added (polyclonal rabbit antibodies ordered from GenScript). The antibody used for each blot is indicated in the figure. Both antibodies were added at a 1:2000 dilution. Blots were incubated with primary for 1 hour at room temperature, rinsed 3x with TBST, then incubated with secondary (Goat-HRP anti-Rabbit IgG) at 1:10000 dilution in TBST (1% milk) for 1 hour. Blots were rinsed 3x with TBST for 15 minutes before adding 12 ml of BioRad Clarity Western ECL Substrate, incubating for 5 minutes, and imaging.

