## Supplementary Information for "Conserved and repetitive motifs in an intrinsically disordered protein drive α-carboxysome assembly"

Turnšek *et al.*, 2023

Supporting Information

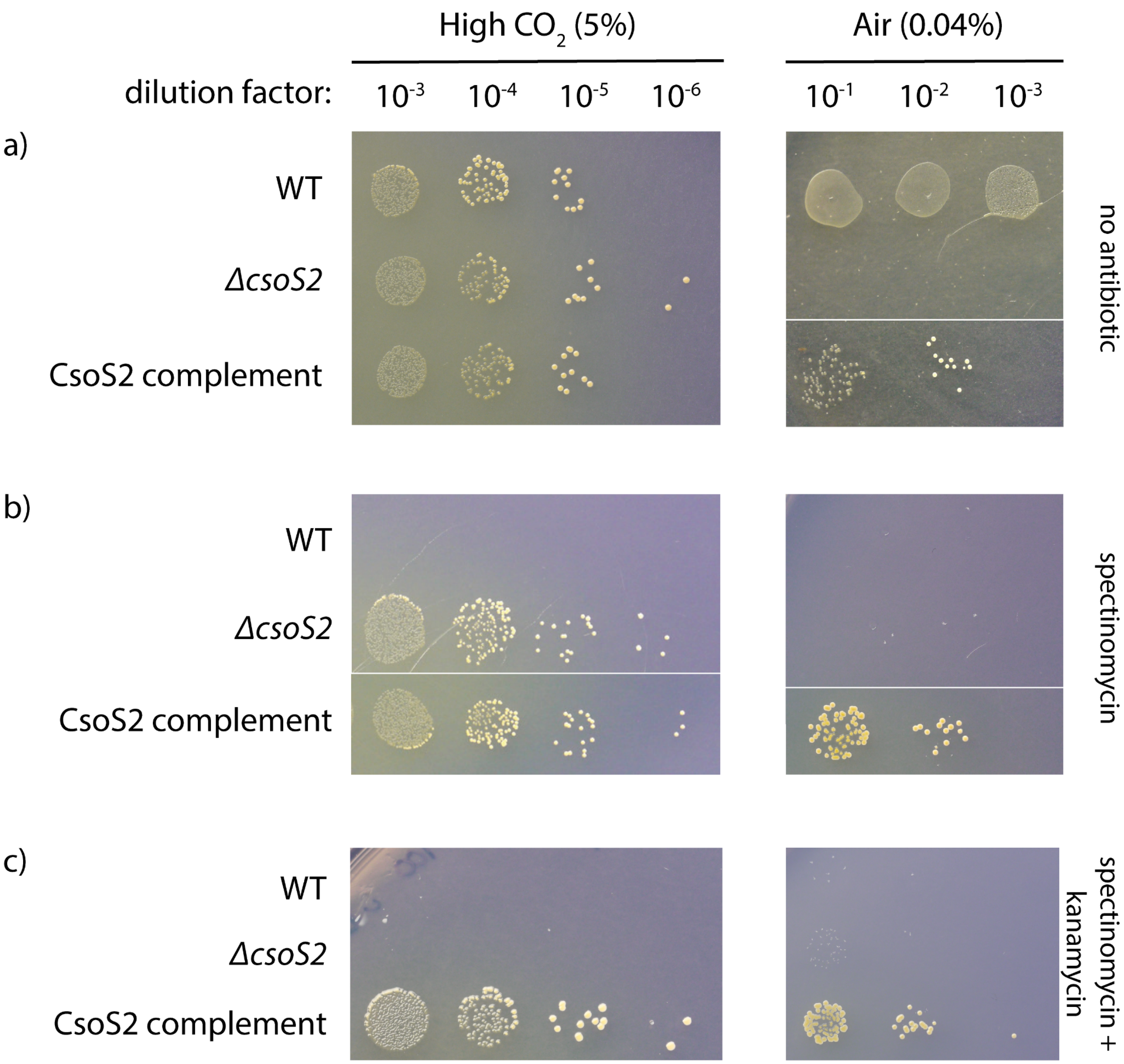


**Figure S1. CsoS2 knockouts are partially complemented in air.** The Δ*csoS2* strain is a knockout of CsoS2 via spectinomycin ORF insertion. The CsoS2 complement strain contains a genomic insertion of IPTG-inducible CsoS2 and kanamycin targeted to a neutral site on the genome in the Δ*csoS2* background strain. WT, Δ*csoS2*, and CsoS2 complement strains were plated in a serial dilution and grown in high CO2 (5% CO2) or air (0.04% CO2) on DSMZ68-agar plates + 100 μM IPTG with (A) no antibiotic, (B) 10 μg/ml spectinomycin, and (C) 10 μg/ml spectinomycin + 2 μg/ml kanamycin. Some images are a composite from different rows on the same plate that were not immediately next to each other, as marked by a thin white line.

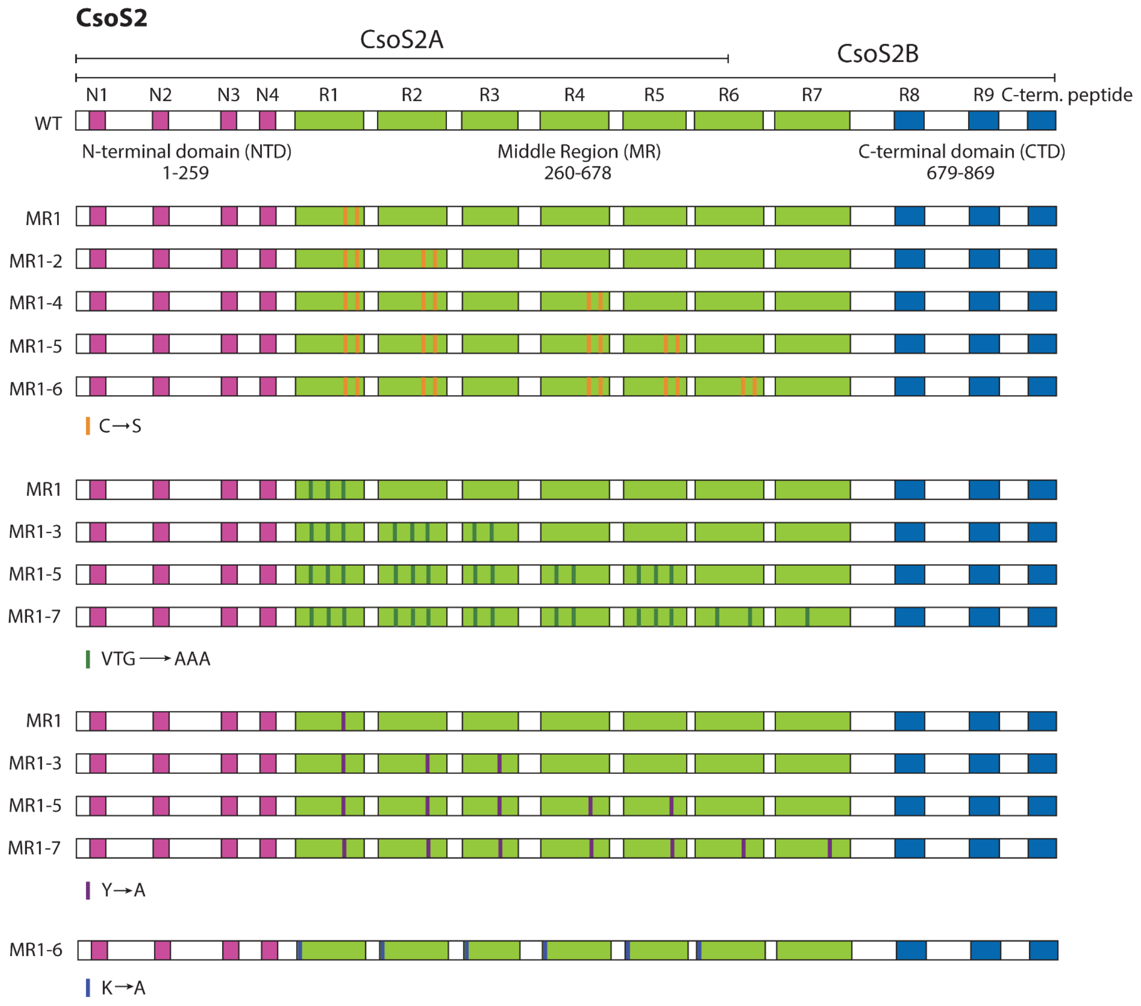


**Figure S2. Illustration of all CsoS2 variants expressed in vivo in *H. neapolitanus*.** Note for the C→S variants, there are no cysteines in R3. Note for the VTG variants, only VTG and VSG sites were mutated.

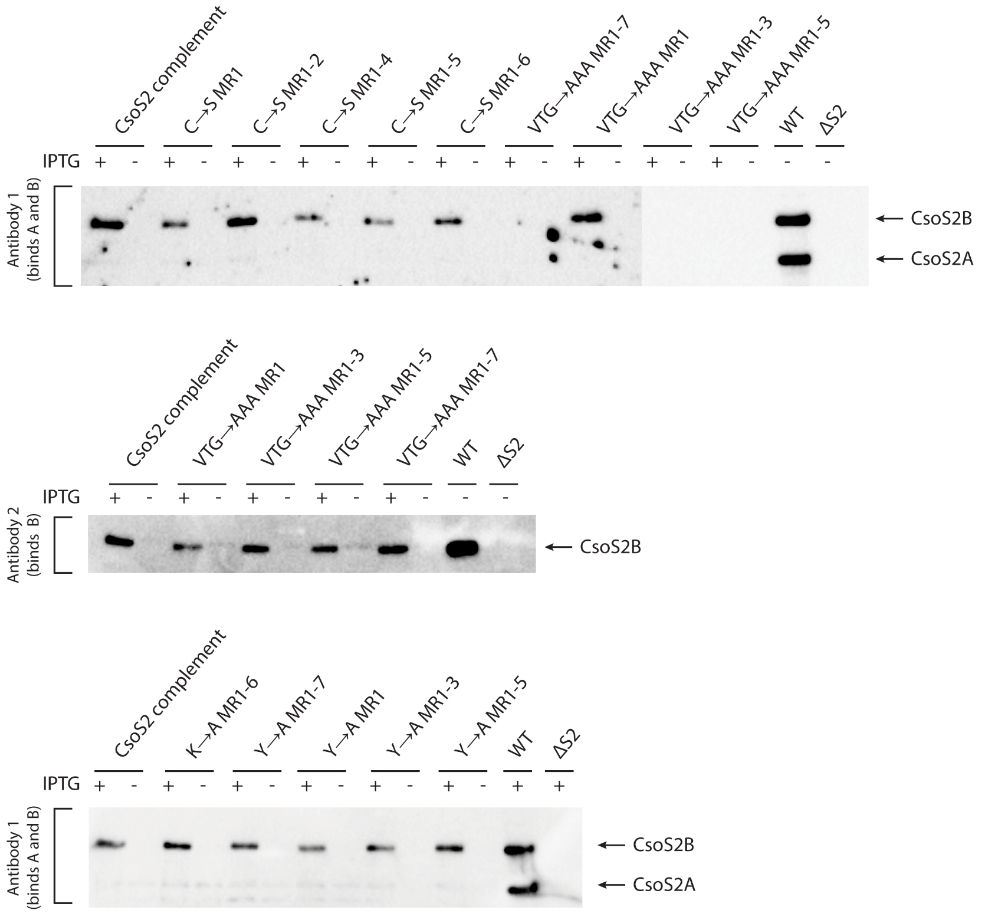


D

| Anti-CsoS2 antibodies | Peptide epitope | Reacts with |
|---|---|---|
| Antibody 1 | CGTEVSRTEKVTGNE | CsoS2A and CsoS2B |
| Antibody 2 | CANRNVPKPEKPGSK | CsoS2B |

**Figure S3. Western blots of complemented CsoS2 and mutant strains.** Strains were grown in liquid culture at high $CO_2$ (5%) +/- 100 μM IPTG and appropriate antibiotics. (A) C→S and VTG→AAA mutants blotted with antibody 1; loading was normalized to ~25 μg/sample. Some antibodies do not bind due to the mutated VTG sequence in the strain. (B) VTG→AAA mutants blotted with antibody 2; loading was normalized to 50 μg/sample. WT was loaded at 12.5 μg to reduce underexposure. (C) K→A and Y→A mutants blotted with antibody 1; loading was normalized to 25 μg/sample. (D) Table of antibodies and their binding epitopes with VTG underlined.

A

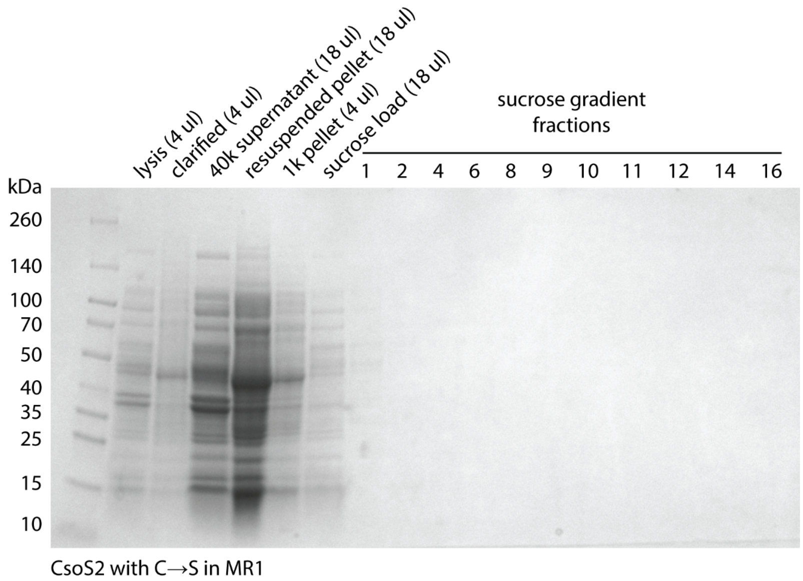


B

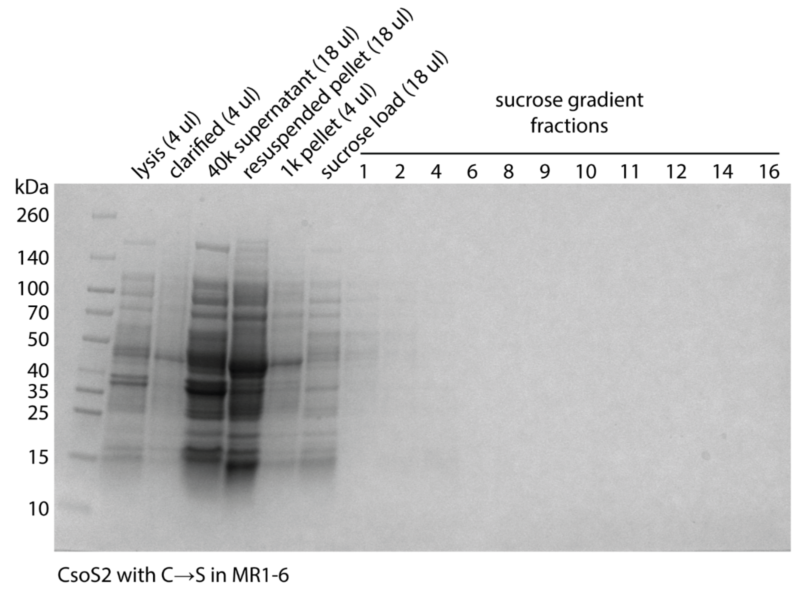


**Figure S4. Carboxysomes with C→S mutations cannot be purified from *E. coli*.** The indicated CsoS2 C→S mutants were expressed in *E. coli* from a plasmid containing the *H. neapolitanus* carboxysome operon and purified using standard methods (see Experimental Procedures for details). (A) PAGE gel of the carboxysome purification with the CsoS2 variant C→S in MR1, and of (B) the CsoS2 variant C→S in MR1-6. Carboxysome bands would be expected to appear around fractions 9-12.

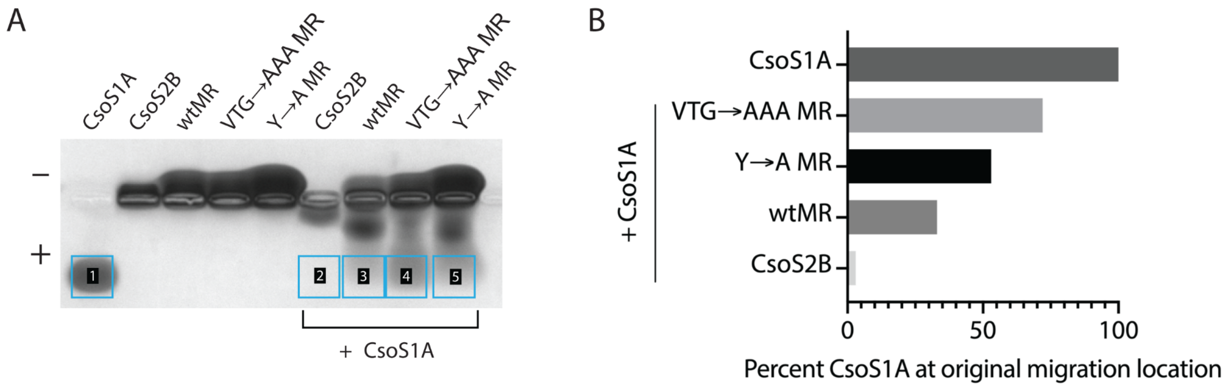


**Figure S5. CsoS2B and wtMR bind to shell protein CsoS1A, and mutated MR variants attenuate binding in a native agarose gel.** (A) Same agarose gel as in Figure 3B, with boxes drawn around quantified areas. (B) Percent of CsoS1A at the original migration location in (A).

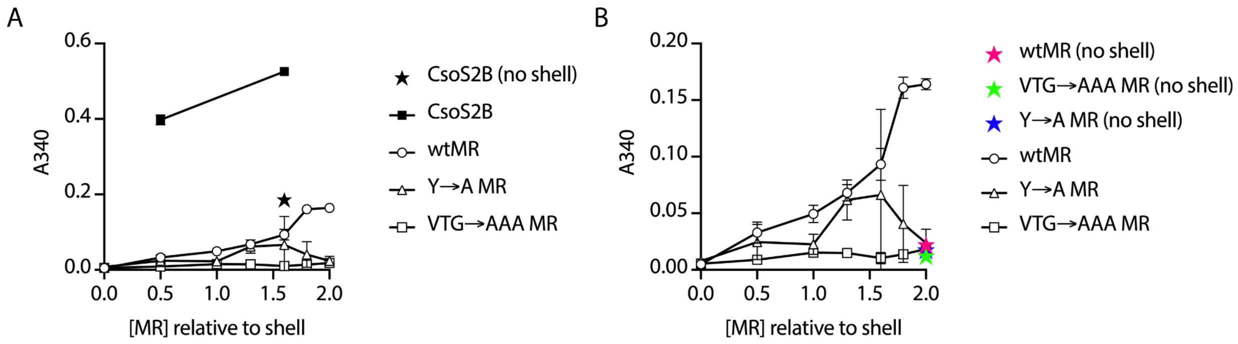


**Figure S6. Shell protein CsoS1A binds CsoS2 with high affinity.** Turbidity assay at 10 minutes of the indicated constructs with CsoS1A. Data are the same as in Figure 3C, but with CsoS2B shown for comparison (A) and no shell controls (A and B). Unobservable error bars are smaller than the datapoint icon.

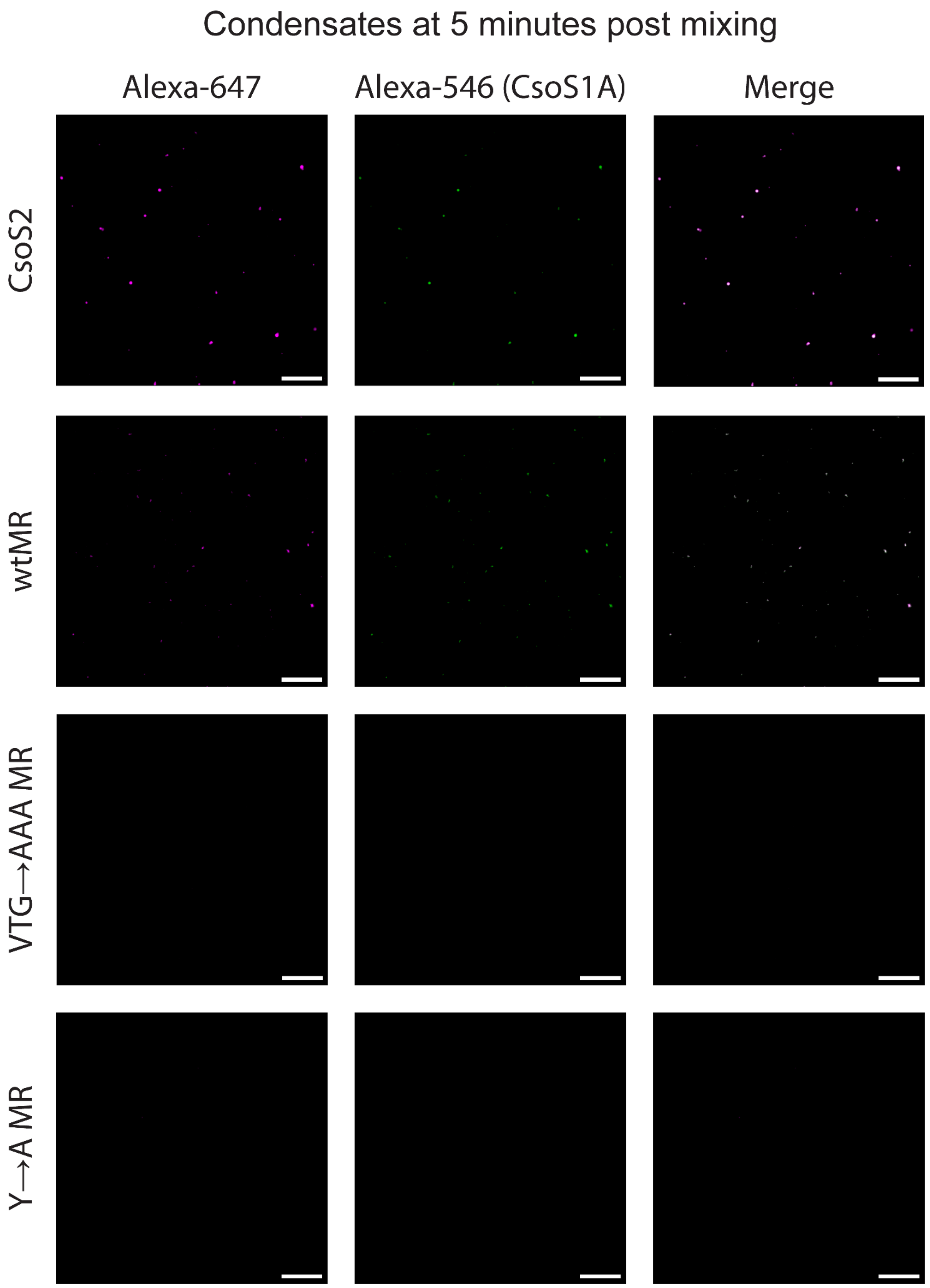


**Figure S7. Smaller condensates at 5 minutes after mixing.** Fluorescence microscopy of the indicated CsoS2 / MR protein variants with added CsoS1A, imaged at 5 minutes post mixing. All CsoS2 / MR variants are labeled in pink, CsoS1A is labeled in green, and the merge appears white at equally overlapping intensities. Scale bar is 20 um.

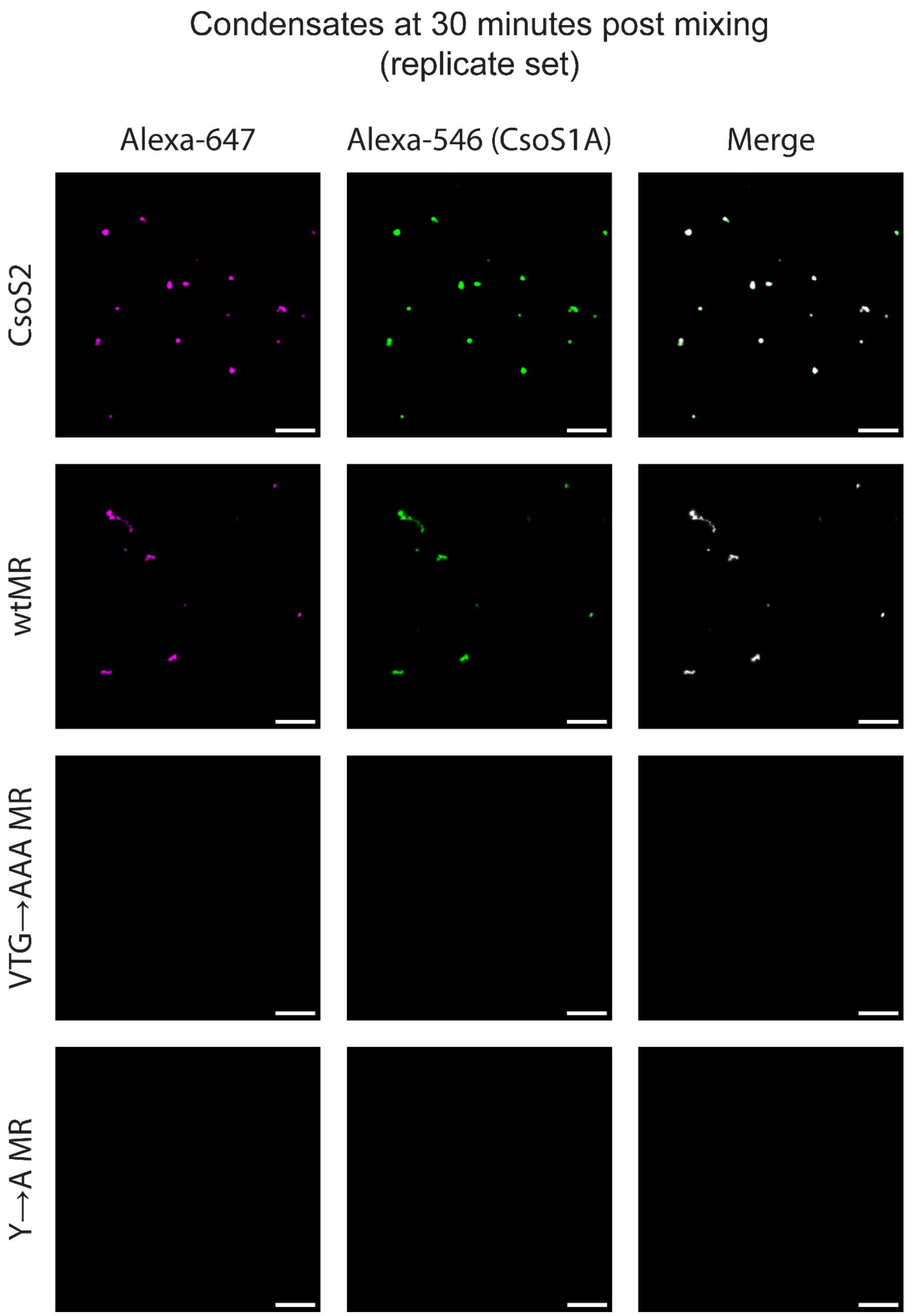


**Figure S8. Larger condensates at 30 minutes after mixing (replicate set).** Replicate set of micrographs in Figure 4 of the indicated CsoS2 / MR protein variants with added CsoS1A, imaged at 30 minutes post mixing. All CsoS2 / MR variants are labeled in pink, CsoS1A is labeled in green, and the merge appears white at equally overlapping intensities. Scale bar is 20 um.

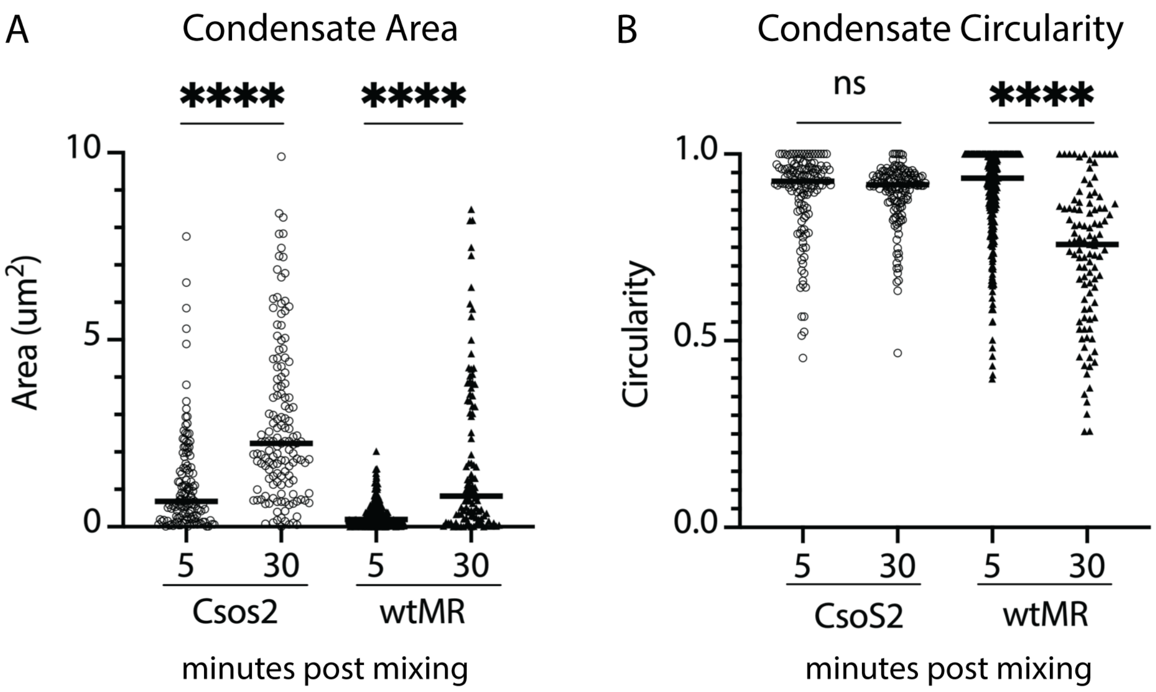


**Figure S9. Over 30 minutes condensate area increases for both CsoS2 and wtMR, but condensates develop into different shapes.** (A) Condensate area in $um^2$ at 5 minutes and 30 minutes post mixing with CsoS1A for both CsoS2 and wtMR. (B) Condensate circularity at 5 minutes and 30 minutes post mixing with CsoS1A for both CsoS2 and wtMR. Circularity is calculated as $4\pi$*area/perimeter$^2$, with 1.0 being a perfect circle and lower values indicating increasing shape elongation. The median is indicated by a black line. Each individual droplet appears as a dot on the plot. Significance of **** is $P \leq 0.0001$ in an unpaired t-test.

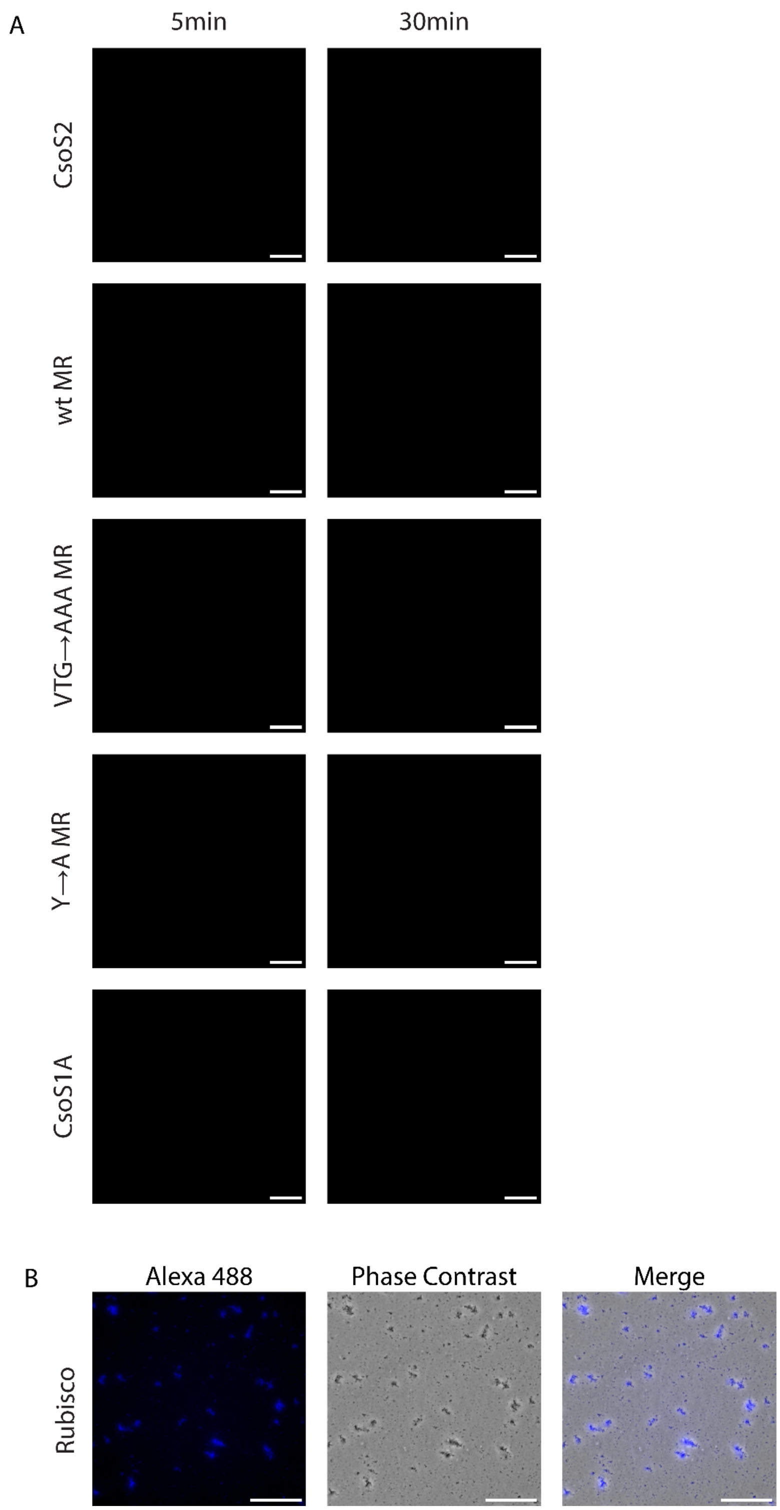


**Figure S10. Individual protein controls show no condensate formation.** (A) Each indicated protein construct was imaged at 5 minutes and 30 minutes. CsoS2 and MR variants were imaged on the Alexa-647 channel. CsoS1A was imaged on the Alexa-546 channel. All scale bars are 20 um. (B) Rubisco imaged at 5 minutes on the Alexa-488 channel and phase contrast, and an image merge. Scale bar is 20 um.

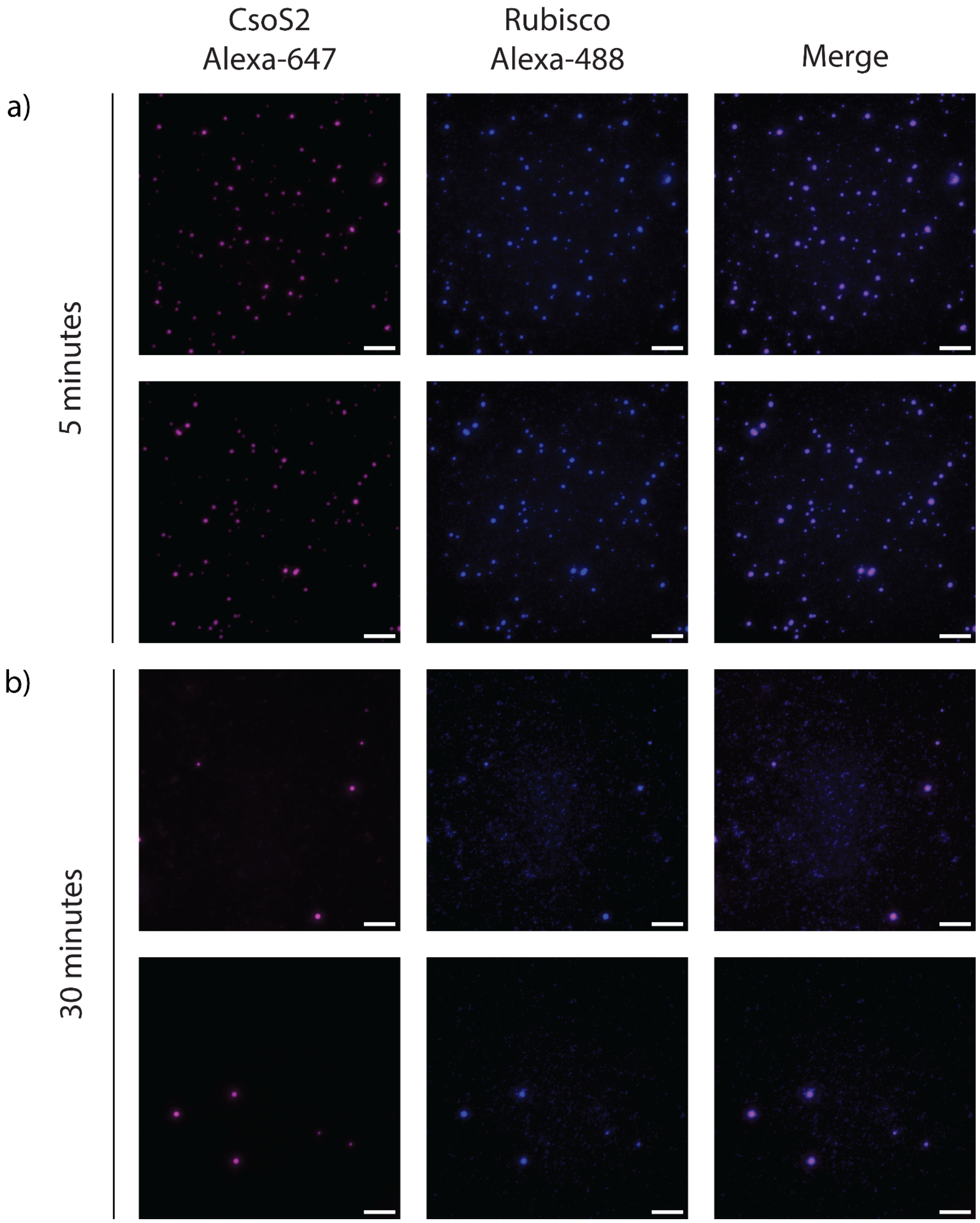


**Figure S11. Rubisco and CsoS2 form condensates that dissociate over time.** CsoS2 is labeled in pink, Rubisco in blue, and the merge is shown in purple. (A) 5 minutes post mixing, two replicate sets, (B) 30 minutes post mixing, two replicate sets. Scale bar is 10 um.

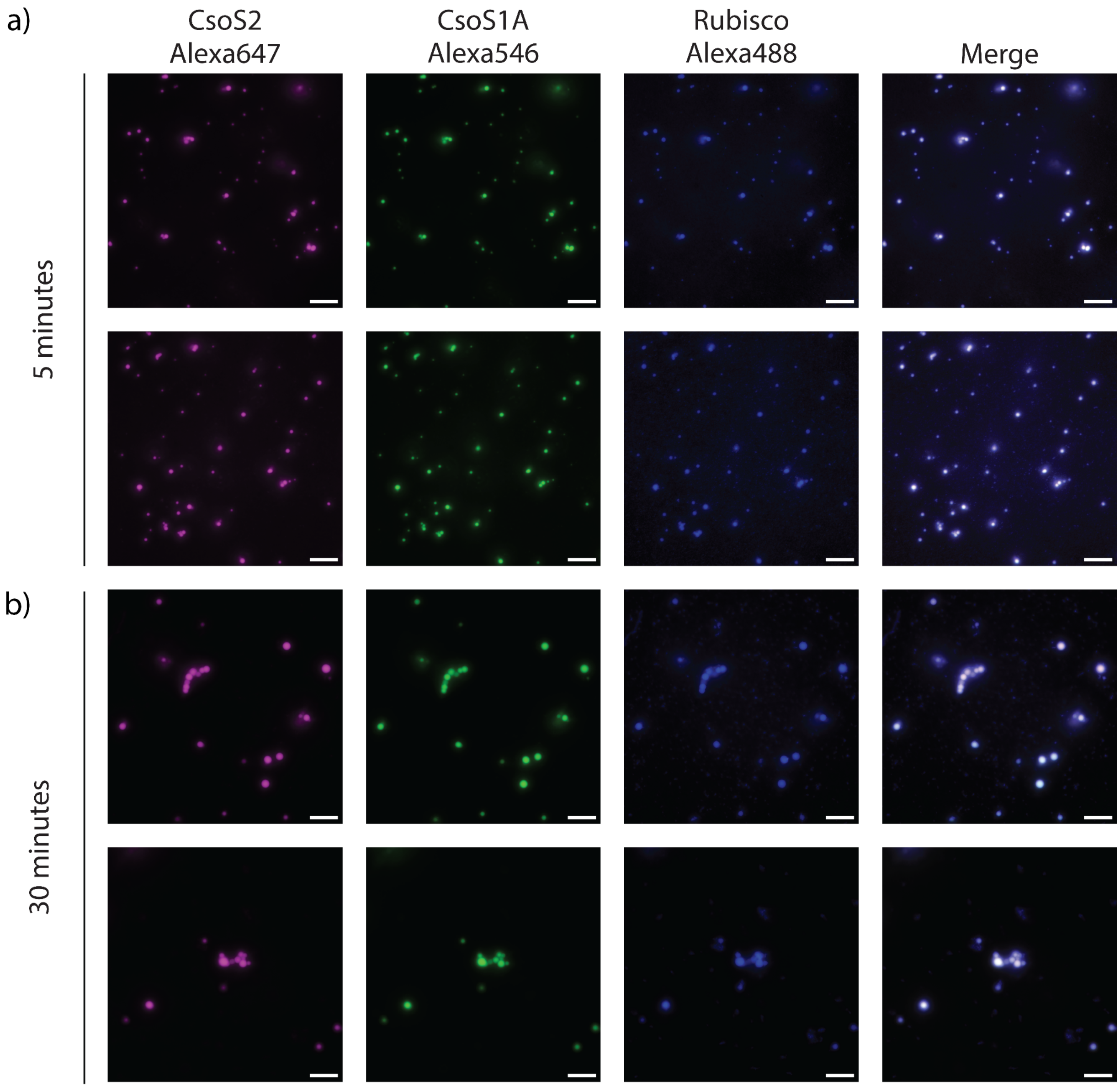


**Figure S12. Rubisco, CsoS2, and CsoS1A form robust spherical condensates that grow in size over 30 minutes.** CsoS2 is labeled in pink, CsoS1A in green, Rubisco in blue, and the merge appears as white. (A) 5 minutes post mixing, 2 replicate sets. (B) 30 minutes post mixing, 2 replicate sets. Scale bar is 10 um.

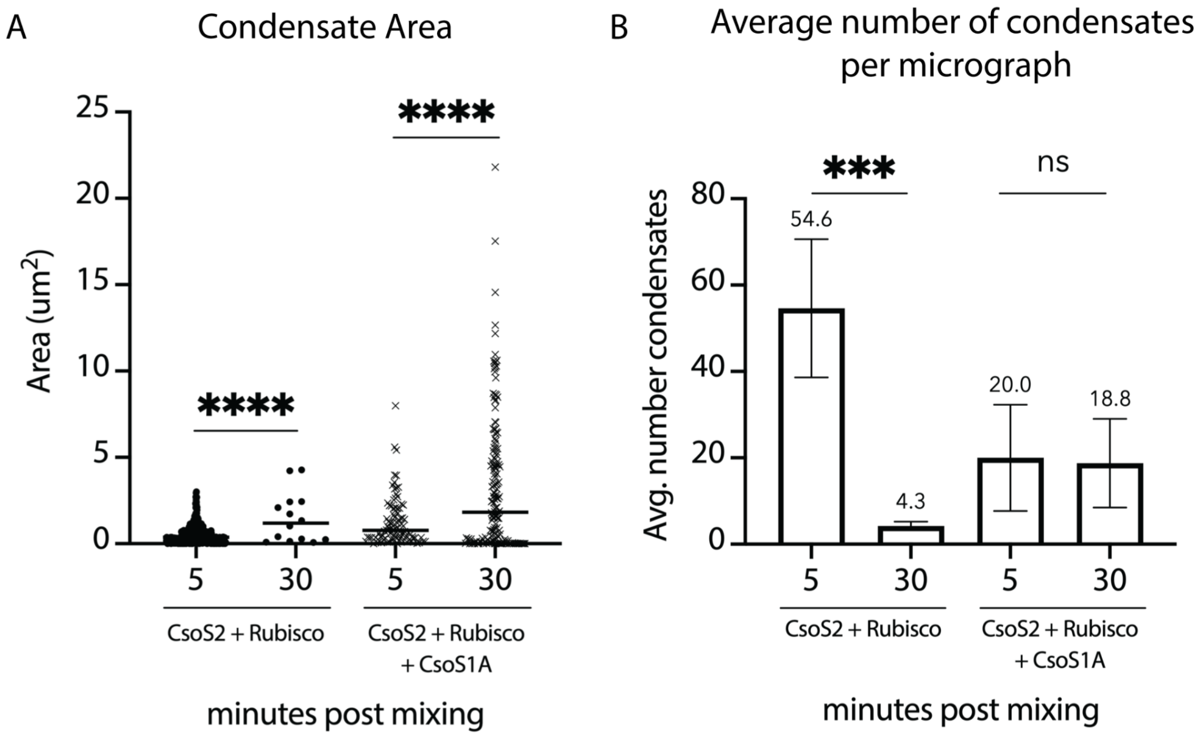


**Figure S13. Addition of CsoS1A leads to an increase in droplet size.** (A) Average area in $um^2$ of CsoS2 + Rubisco condensates and CsoS2 + Rubisco + CsoS1A condensates at 5 minutes and 30 minutes post mixing. The median is indicated by a black line. Each individual droplet appears as a dot on the plot. Significance of **** is P ≤ 0.0001 in an unpaired t-test. (B) Average number of condensates per micrograph, which measured 83.2 x 83.2 um. The average is written above each bar. Significance of *** is P ≤ 0.001 in an unpaired t-test. Ns, no significance.

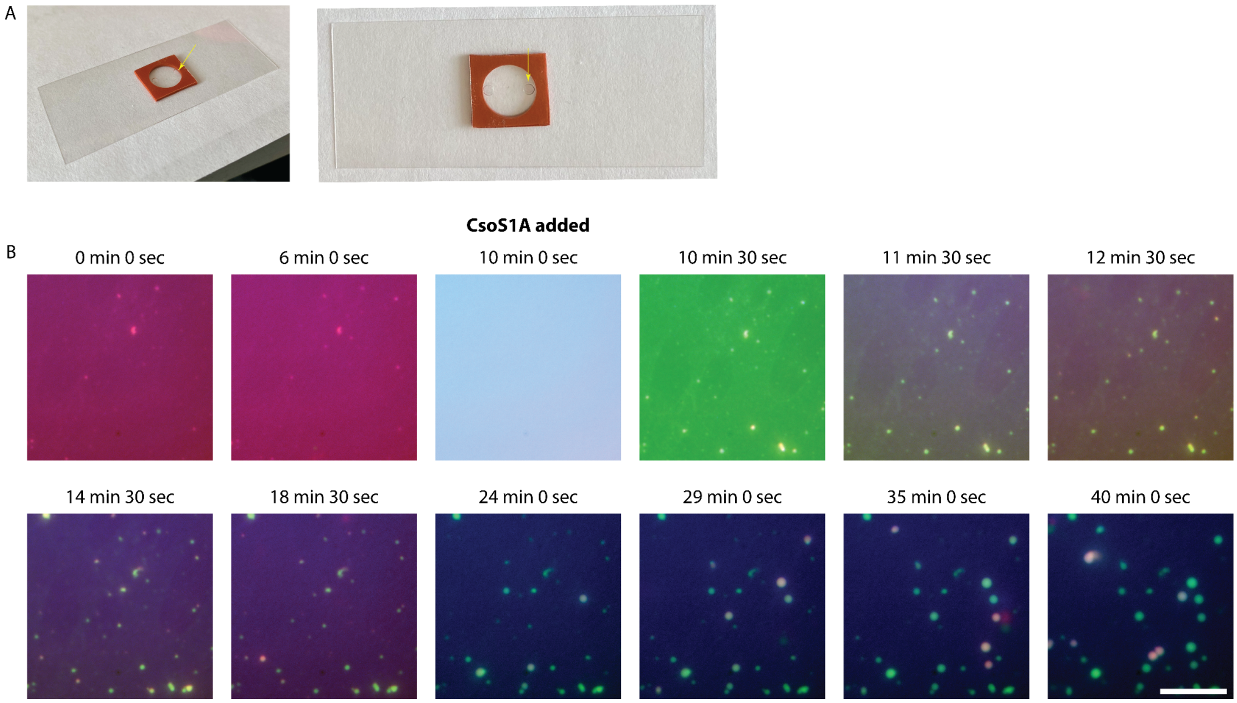


**Figure S14. Adding CsoS1A to Rubisco + CsoS2 nucleates condensate formation.** (A) Gasket setup; the gasket was affixed to a microscope slide, and protein solution was added with a pipette into the pore indicated by the yellow arrow. (B) Rubisco (blue) and CsoS2 (red) were premixed and added to a gasket affixed to a microscope slide. At 10 minutes, CsoS1A (green) was added. Droplets appear in the focus plane as they condense and adhere to the slide. Times were rounded to the nearest 30 seconds. Scale bar is 10 um.

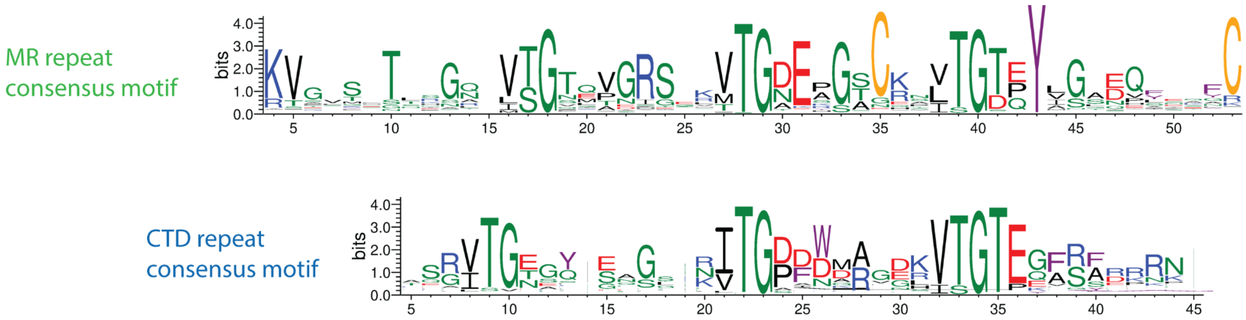


**Figure S15. CTD repeat consensus motif**. Comparison between the MR repeat consensus motif and the CTD repeat consensus motif. Sequence logo of the MR repeat was generated from an alignment of 1662 MR repeats identified across 272 dereplicated CsoS2 sequences. Sequence logo of the CTD repeat was generated from an alignment of 528 CTD repeats identified across the same 272 dereplicated CsoS2 sequences. Blue is basic, red is acidic, green is polar/small, black is hydrophobic, yellow is cysteine, purple is aromatic.
